# Protein concentrates from pilot-scale biorefining of clover grass: Interfacial properties and their effects on the physical and oxidative stability of fish oil-in-water emulsion

**DOI:** 10.64898/2026.02.06.704303

**Authors:** Narjes Badfar, Mette Lübeck, Charlotte Jacobsen, Simon Gregersen Echers

## Abstract

Clover grass blends are promising sources of nutritional and techno-functional proteins, but currently mainly utilized for animal feeding. The application as a physical and oxidative stabilizer in food emulsions remains underexplored. In this study, the stabilizing effects of clover grass proteins (CGPs), produced through a pilot-scale, two-stage membrane filtration process yielding a native GPC concentrate (DC), as well as enzymatic hydrolysate hereof (DCH), were compared with commercial plant proteins (soy and pea) and animal sources (sodium caseinate). Both DC and DCH produced emulsions (0.4% (w/w) protein and 5% fish oil) with smaller size droplets and larger electrostatic repulsion between droplets compared to the other proteins tested. Moreover, DC and DCH exhibited higher protection against the generation of both primary and secondary oxidation products. Furthermore, emulsions stabilized with CGPs were well-protected from off-flavor compounds. Mass spectrometry-based proteomics analysis revealed that DC included a high RuBisCO content (38%) and the membrane process successfully depleted pigment-binding proteins affiliated with grassy color and sensory attributes. Moreover, DC was enriched (compared to the initial green juice) in known antioxidant proteins, constituting 10% of the total protein. In the hydrolysate (DCH), 30% of the total MS1 peptide signal originated from peptides predicted as probable free radical scavengers. These findings demonstrate that refined, native CGP, as well as its hydrolysate, improved both physical and oxidative stability of emulsions compared to plant and animal-based reference proteins due to a high endogenous antioxidant properties of the protein.

## 1. Introduction

The application of synthetic emulsifiers in the food industry might cause health problems and toxic symptoms for consumers following prolonged use (Dammak et al., 2020). As such, proteins and protein-based ingredients are gathering attention for stabilizing food emulsions by creating highly elastic networks at the droplet surface. Due to their amphiphilic nature, some proteins can migrate to the interface and decrease the interfacial tension in emulsion systems (Mileti et al., 2023). Therefore, utilizing natural emulsifiers in different industries, especially the food sector, is highly desirable. Alternative proteins, particularly of plant origin, are being considered as replacements for animal-based proteins such as caseins, lactoglobulins, and albumins, due to their higher sustainability profile as well as potential positive health impacts, such as glycemic reduction and antitumor properties (Mileti et al., 2023).

Among plant-based proteins, green leafy biomass such as grasses are promising sources of alternative proteins for food applications (Pérez-Vila et al., 2024). However, proteins from blends of clovers and grasses (clover grass) and alfalfa are conventionally utilized as animal feed, priced below food proteins (Pearce & Brunke, 2023; Santamaría-Fernández & Lübeck, 2020). Nevertheless, because green leaves are rich in the enzyme ribulose-1,5-bisphosphate carboxylase/oxygenase (RuBisCO), they have been proposed as a potential source of food-grade protein for human consumption several decades ago (Barbeau & Kinsella, 1988). RuBisCO is recognized as a superior source of protein among other plant protein sources, owing to reasons attributed to sulfur-containing amino acids such as cysteine and a generally well-balanced amino acid profile for human consumption, but still remains an underutilized source of protein in food applications (Pérez-Vila et al., 2024). Denmark is well-known for cultivating grasses and possesses a share of 40% of the EU grass seeds production, especially clover grass blends (Santamaría-Fernández & Lübeck, 2020). This is due to Denmark’s appropriate climate for producing various types of crops. Conventional methods for extracting leaf proteins for feed, such as heat treatment, pH adjustment, or lactic acid fermentation, are unsuitable for human consumption due to their production of a bitter taste, relatively low protein content (35–50%), and the co-extraction of undesirable phytochemicals (Santamaría-Fernández & Lübeck, 2020). To address these limitations, a mild membrane separation process that effectively removes chlorophyll and the bitter taste meanwhile producing a highly soluble and RuBisCO-rich food-grade protein product has recently been developed (Gregersen Echers et al., 2025; Mattsson et al., 2025).

Fish oil is commonly utilized to produce oil-in-water emulsions. Beyond its beneficial effects on health conditions, attributed to the high content of polyunsaturated fatty acids (PUFAs), fish oil is widely recognized as a well-known model for investigating oxidation stability in functional food products. The presence of PUFAs makes fish oil highly susceptible to oxidation, which has adverse effects on the nutritional value and sensory properties, such as changing the color, odor, flavor, and texture of the final product (García-Moreno et al., 2016). Applying proteins as emulsifiers at the oil-water interface influences the rheological properties of the emulsion system. An interfacial layer with a viscoelastic behavior and ability to endure the expansion is considered a stronger stabilizer of an interfacial layer (Mileti et al., 2022). Several studies have shown that plant proteins can improve the physical and oxidative stability of emulsions (García-Moreno et al., 2020; Pérez-Gálvez et al., 2024; Pérez-Vila et al., 2024; Tan et al., 2022; Yesiltas et al., 2022) Despite the potential of clover grass proteins (CGPs) as a functional protein ingredient (Badfar et al., 2026; Echeverria-Jaramillo et al., 2025, 2026), there remains a significant knowledge gap regarding the oxidative and physical stability when used as an emulsifier. As such, dilatational rheology plays a vital role in delving into the CGP ability and potential application as a stabilizer in food emulsion systems.

This study primarily focused on the interfacial properties of CGP and its hydrolysate in a medium-chain triglyceride (MCT) oil phase, providing a dynamic and molecular view of the protein behavior at the interface between MCT oil and water. These findings were further linked to the application of CGPs in a fish oil-in-water emulsion system to investigate their physical and oxidative characteristics using a wide range of methods. Moreover, three control samples were used for comparison plant-based proteins (pea protein isolate and soy protein isolate) and an animal-based control (sodium caseinate). This allowed the properties of CGP and its hydrolysate to be compared with these well-known and commercially used proteins. Furthermore, using mass-spectrometry-based proteomics, we investigated the protein- and peptide-level composition of the protein and hydrolysate, respectively, to obtain insight into what proteins and peptides may attribute to the observed functionality. Ultimately, this study provides both novel insight on the use of CGP and hydrolysates as emulsifiers meanwhile linking bulk functionality to molecular composition.

## 2. Material and methods

### 2.1. Materials and sample preparation

This study used a commercial blend of clover grass (16% red clover, 10% white clover and 74% perennial grasses (including perennial ryegrass, fescue, and hybrids) to produce two different clover grass protein (CGP) samples. The biomass was grown at Ausumgaard, Denmark (56.4201° N, 8.6084° E) and was harvested on August 17^th^ 2023. Green juice, obtained by screw pressing freshly harvested biomass, was produced using a two-stage membrane filtration, as previously described (Mattsson et al., 2025). In contrast to our previous study (Badfar et al., 2026) using a demonstrator-scale process (∼15-20kg clover grass), the CGP in this work was produced in a pilot-scaled version (∼600kg clover grass) of the same process (Lübeck et al., 2025). Following initial removal of “green protein” in the first stage filtration (1330 L total feed volume), the retentate from the second-stage filtration was subsequently subjected to diafiltration to obtain a purified protein concentrate (DC), of which a part was lyophylized (Telstar LyoQuest freeze dryer, Azbil Co., JP). Another part of the DC, was hydrolyzed using 1% (w/w) Formea® Prime (trypsin) (140 KMTU/g), purchased from Novonesis (Bagsværd, Denmark), to produce a hydrolysate (DCH). The hydrolysis was performed as free-fall pH hydrolysis initiated at the optimum conditions of the enzyme (37 °C and pH 8) in a shaking water bath with a constant condition. The hydrolysis process was conducted for 2.75 h based on previous functionality-based optimization screening (Badfar et al., 2026). The mixture subsequently was heated up to 90 °C for 15 min for trypsin inactivation. After inactivation, the mixture was centrifuged at 2000 ×g at 22 ^°^C for 10 min. The supernatant was collected and lyophylized. Pea protein isolate (PPI) (Emsland group, Emlichheim, Germany), soy protein isolate (SPI) (Bulk, Essex, UK), and sodium caseinate (Na-Cas) (Arla Foods, Viby J, Denmark) were used as references. For proteomics analyses, the initial feed stream for the two-stage membrane filtration (i.e. the green juice) was included as reference.

### 2.2. Crude protein content and degree of hydrolysis (DH%)

The crude protein (CP) content of the clover grass samples was analyzed using the Dumas method (Elemental Analysensysteme GmbH, Germany). The measurements were carried out based on the total nitrogen content, and the results were converted to CP content using a factor of 6.25 (n=3).

The degree of hydrolysis was measured based on the percentage of peptide bond cleavage in the liquid hydrolysate using the OPA (O-Phthaldialdehyde) assay, according to (Bjørlie et al., 2024).

### 2.3. Interfacial properties

#### 2.3.1. Interfacial Tension (IFT) and Dilatational Rheology

Interfacial properties of CGPs (DC, DCH) and control samples (PPI, SPI, and Na-Cas) were assessed using interfacial tension measurement and dilatational rheology tests with the pendant drop method using a drop tensiometer (OCA20, Dataphysics GmbH, Germany). A droplet of 0.1% (w/v) protein solution with a size of 35 μL was shaped at the edge of the needle (d=1.83 mm) in the quartz-glass cuvette containing medium-chain triglyceride (MCT) oil. The protein solution’s interfacial tension (IFT) at the oil-water interface was recorded for 30 min at room temperature, and the droplet volume was controlled and constant to balance out and reach equilibrium. The IFT measurement was carried out based on the Young-Laplace equation and dilatational rheology was performed as previously described (Badfar et al., 2025). To test the dilatational rheology, following an equilibration time of 3 h, the linear viscoelastic regime was determined. The deformation amplitude was altered manually (1.00%, 2.75%, 4.50%, 6.25%, and 8.00% interfacial area) at a constant frequency of 0.01 Hz, where each oscillation involved 5 cycles, and, before each step, the interfacial film was allowed to recover for 90 s. The frequency sweeps were performed after the amplitude sweeps, where the frequency differed from 0.01 to 0.10 Hz at an amplitude of 4.00%.

### 2.4. Emulsion properties

To produce the 5% (w/w) oil-in-water emulsion, 0.4 % (w/w, protein basis) CGP (DC or DCH) and control samples (PPI, SPI, or Na-CAS) were dissolved in acetate-imidazole buffer (pH=7) in a water bath shaker (130 rpm, 50 °C, 2 h). The dispersed samples were hydrated at 130 rpm overnight in the dark at RT. The next day, homogenization was conducted in two steps. First, the protein solution was homogenized by ultra turrax (IKA Werke GmbH &. Co., Staufen, Germany), and cod liver oil (Vesteraalen, Sortland, Norway) was gradually introduced to the water phase during the first minute of homogenization at 16,000 rpm. The first step of homogenization lasted 60 s to form a pre-emulsion. Subsequently, a second homogenization step was conducted using a high-pressure homogenizer (Panda Plus 2000, GEA Niro Soavi, Lübeck, Germany) under a constant temperature below °26 C (9 Pas with three passes).

### 2.5. Storage experiment and sampling

After emulsion preparation, the emulsions were sorted in the 30 ml dark bottles for the physical stability experiments. The physical stability of the emulsions was evaluated by ζ-potential, droplet size determination, and visual assessment of the emulsions during the 8-day storage period.

To determine oxidative stability, emulsions were immediately aliquoted into 30 ml dark bottles and divided based on the experiments and aq. FeSO4 (80 mM) was added to accelerate lipid oxidation. The oxidative stability tests were performed during a 9-day storage period. At each sampling point, the containers were flushed with nitrogen gas and sealed with plastic caps, transferred to a −40 °C freezer, and stored until further analysis.

### 2.6. Physical stability of the emulsions

#### 2.6.1. ζ-potential determination of emulsion droplets

The ζ-potential analysis was conducted on all emulsion samples on days 1 and 8 using a ZETASIZER NANO ZS along with a DTS1070 cell (Malvern Instruments Ltd., Worcestershire, UK) at RT. Ten μL of the emulsion samples were diluted in 5 ml imidazole buffer (pH=7), and zeta potential was measured with 100 runs for each emulsion sample (n=3).

#### 2.6.2. Emulsions droplets size measurement (D[4,3] and D[3,2])

The droplet size measurements were performed using laser diffraction in a MASTERSIZER 2000 (Malvern Instruments, Ltd., Worcestershire, UK). The volume (D[4,3]) and surface (D[3,2]) mean diameter measurements were carried out on day 1, 3, and 8 of the storage period (n=3).

#### 2.6.3. Visual assessment of emulsion

The emulsions’ appearance was assessed visually, and the oil-rich phase accumulation was determined as the oil index (OI) and calculated as:

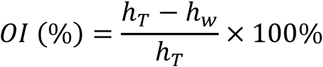

Where h_T_ is the total height of the emulsion and h_w_ is the height of clear water.

### 2.7. Oxidative stability of the emulsions

#### 2.7.1. Peroxide value (PV)

The primary lipid oxidation products formed during oil-water emulsion ageing were assessed by determination of the peroxide value (PV). Lipids were extracted from emulsion samples according to the method of Bligh & Dyer, (1959), using chloroform/methanol/water (1:1:0.5, w/w). Each extraction was performed in duplicate. The peroxide value was determined on the lipid extracts using the ferric thiocyanate method described by Shantha & Decker (1994). Briefly, lipid hydroperoxides oxidize ferrous ions (Fe²⁺) to ferric ions (Fe³⁺), which subsequently react with thiocyanate to form a red ferric-thiocyanate complex. The absorbance of this complex was measured spectrophotometrically at 500 nm (Shimadzu UV-1280, Holm & Halby, Brøndby, Denmark). Peroxide values were calculated from the absorbance measurements and expressed as kg milli equivalent O₂/kg oil. Measurements were conducted in duplicate for each lipid extract.

#### 2.7.2. Tocopherols consumption measurement using HPLC

Tocopherol analysis was carried out by high-performance liquid chromatography using an Agilent 1100 system (Agilent Technologies, Palo Alto, CA, USA) coupled to a fluorescence detector. Lipids obtained from the Bligh and Dyer extraction were used for analysis. Two g of lipid extract was evaporated under a gentle nitrogen stream and subsequently reconstituted in 10 mL of n-heptane. Aliquots (1 mL) of the resulting solution were transferred into individual vials prior to chromatographic analysis. Separation was achieved under isocratic conditions using an n-heptane/2-propanol mobile phase (100:0.4, v/v) at a flow rate of 1 mL/min. A 20 µL sample volume was injected onto a silica column (Waters Spherisorb, 3 µm, 4.6 mm × 150 mm), fitted with a matching silica guard column (Waters Spherisorb, 5 µm, 4.6 mm × 10 mm). Tocopherols were detected by fluorescence with excitation and emission wavelengths set at 290 nm and 330 nm, respectively, in accordance with the AOCS method (1998). Quantification was performed using an external calibration curve generated from a tocopherol standard mixture containing α-, β-, γ-, and δ-tocopherols (Calbiochem 613424). All analyses were conducted duplicate, and tocopherol concentrations were calculated based on authentic standards.

#### 2.7.3. Secondary Volatile Oxidation products by headspace GC-MS

Volatile oxidation products were analyzed using an automated dynamic headspace system coupled to gas chromatography–mass spectrometry (GC–MS) (Gerstel GmbH & Co. KG., Mülheim an der Ruhr, Germany). Approximately 1 g of emulsion sample was weighed into a 10 mL headspace vial and supplemented with 30 mg of an internal standard solution containing 4-methyl-1-pentanol (30 µg/g in rapeseed oil). Samples were prepared and analyzed in triplicate. Headspace sampling was performed automatically according to a programmed sequence: The sealed vials were incubated at 60 °C for 4 min with agitation at 300 rpm, applying occasional mixing cycles (10 s on, 1 s off). Volatile compounds were subsequently purged from the vial headspace using nitrogen at a flow rate of 50 mL/min for 20 min and collected on thermal desorption tubes packed with Tenax GR 300 (Gerstel GmbH & Co. KG., Mülheim an der Ruhr, Germany) sorbent. Residual moisture was removed from the sorbent tubes by applying a nitrogen purge of 30 mL/min for 3 min. Trapped volatiles were thermally desorbed and transferred to the GC–MS system via a cooled injection system (CIS 4). Desorption was initiated at 40 °C, followed by rapid heating to 280 °C (720 °C/min) and holding for 5 min. The analytes were then introduced into an Agilent 6890 N gas chromatograph (Agilent Technologies, Santa Clara, USA) coupled to a 5973 inert mass-selective detector (MS 5973, Agilent Technologies, Santa Clara, USA). Separation was achieved using a DB-1701 fused silica capillary column (30 m × 0.25 mm I.D., 0.5 µm film thickness; DB 1701, Agilent Technologies, J&W GC Columns, USA). Mass spectrometric detection was performed in electron ionization mode at 70 eV, with spectra recorded over an m/z range of 30–250. The oven temperature program consisted of an initial hold at 35 °C for 3 min, followed by heating to 140 °C at 3 °C/min, to 170 °C at 5 °C/min, and to 240 °C at 10 °C/min, with a final hold of 8 min. Volatile compounds were identified by comparison of mass spectra with reference libraries (Wiley 138 K, John Wiley and Sons, Hewlett-Packard). Quantification was carried out using external calibration curves prepared at seven concentration levels (0.5–250 µg/mL) for selected standards, including (t,t)-2,4-heptadienal, 1-penten-3-ol, c-4-heptenal, octanal, t-2-hexenal, t-2-pentenal, 1-penten-3-one, pyridine, t-2-butenal, butanal, pentanal, and 1-pentanol. Calibration standards were dissolved in rapeseed oil and analyzed under the same conditions as emulsion samples. Results are reported as mean ± standard deviation of triplicate measurements and expressed as ng of volatile compound per g of emulsion.

### 2.8. Proteomic characterization by LC-MS/MS

In addition to the two CGPs, the initial green juice feed stream for membrane filtration (DM 3.25%, CP 16.6%) for the first stage filtration operation (included as reference) was analyzed by LC-MS/MS-based quantitative shotgun bottom-up proteomics (BUP). Samples were prepared for analysis using the iST kit for plant tissue (PreOmics, Germany), as previously described (Badfar et al., 2026; Gregersen Echers et al., 2026). Because no crude biomass was included in this study, the workflow did not include the focused ultrasonication step. Briefly, sample aliquots corresponding to 70 µg protein (liquid for Feed (green juice) and resuspended (aq.) powder for DC and DCH) were mixed with 100 µL LYSE buffer and incubated for 10 minutes at 95°C in a Thermomixer (Eppendorf, Germany) at 1000 rpm for protein solubilization, reduction, and alkylation. After cooling, samples were digested for 2 hours with resuspended trypsin/LysC and washed according to manufacturer guidelines. As DCH is a hydrolysate, this was not further digested but merely added the resuspension buffer with no added enzyme for the digest step. Lastly, clean digests were dried down and resuspended before concentration estimation using A280 (1A = 1mg/mL).

LC-MS/MS analysis was performed as previously described (Badfar et al., 2026; Gregersen Echers et al., 2026). Briefly, chromatography was performed using an EASY nLC-1200 ultra-high-performance liquid chromatography system (Thermo Scientific, USA) equipped with a PEPMAP trap column (75 μm x 2 cm, C18, 3 μm, 100 Å) connected in-line to a reversed-phase PEPMAP analytical column (75 μm x 50 cm, C18, 2 μm, 100 Å) using a step-wise gradient from 5% to 100% buffer B (80% acetonitrile, 0.1% formic acid (VWR, Denmark)) in buffer A (0.1% formic acid (Fisher Scientific, Denmark) over 60 minutes. Following chromatographic separation, peptides were introduced to a Q Exactive HF tandem mass spectrometer (Thermo) via an ESI ion source. Data acquisition was performed as full MS/ddMS2 Top20 DDA in positive mode with an MS1 range of 300-1600 m/z.

LC-MS/MS data was analyzed in MaxQuant (2.2.0.0) using standard setting and searched against all UniProt accensions for *Lollium perenne* (perennial ryegrass, taxid: 4522, 825 entries), *T. repens* (white clover, taxid: 3899, 477 entries) as well as the reference proteomes of *Trifolium pratense* (red clover, UP000236291, 60,146 entries) *Brachypodium distachyon* (purple false brome, UP000008810, 44,786 entries), and *Trifolium subterraneum* (subterranean clover, UP000242715, 36,725 entries) and all entries from the *Festuca* genus (taxid: 4605, 1,498 entries) were also included. All protein lists were downloaded from Uniprot (“UniProt: The Universal Protein Knowledgebase,” 2017) on December 2^nd^ 2024. Protein quantification was performed using both label-free quantification, MaxLFQ (Cox et al., 2014), as well as intensity-based absolute quantification, iBAQ (Schwanhäusser et al., 2011). The mass spectrometry proteomics data have been deposited into the ProteomeXchange Consortium via the PRIDE (Perez-Riverol et al., 2022) partner repository with the dataset identifier PXD070032 and doi 10.6019/PXD070032.

#### 2.8.1. Downstream analysis and bioinformatics using MaxLFQ and iBAQ data

Differential analysis of MaxLFQ quantified data was performed using Mass Dynamics 3.0 (Quaglieri et al., 2022), as previously described (Badfar et al., 2026; Gregersen Echers et al., 2026). Following imputation of missing values, the whole dataset was analyzed for differential proteins by ANOVA and visualized using heatmaps (Euclidean distance = 4) and principal component analysis (PCA).

iBAQ data was used to quantify the relative molar protein distribution within each sample by mean riBAQ within sample triplicates, as previously described (Badfar et al., 2026; Gregersen Echers et al., 2026). Prior to quantitative analysis, common contaminants and false positives were filtered. To evaluate RuBisCO retention and removal of unwanted proteins from the Feed stream, the total riBAQ for the following families/groups was computed across the full dataset: RuBisCO (rbc), Chlorophyll a-b binding protein (CBP), Photosystem I & II proteins (PI/II). In addition, known antioxidant proteins previously identified in substantial abundances and linked to *in vitro* antioxidant activity in *L. perenne* were quantified in a similar way. According to (Danner Aakjaer Pedersen et al., 2025), these include: ferredoxin-NADP reductase (FNR), lactoylglutathione lyase (Glyoxalase I; Glo1), glutathione S-transferase (GST), dehydroascorbate reductase (DHAR), Thioredoxin-dependent peroxiredoxin (TPx), Superoxide dismutase (SOD), L-ascorbate peroxidase (APX), Glutaredoxin-dependent peroxiredoxin (GPx), Peroxidase (Px), and Peroxiredoxin Q-like (ycf33).

As DCH does not contain proteins but only peptides as a hydrolysate, the antioxidant potential of the peptides was evaluated using AnOxPePred (Olsen et al., 2020). All identified peptides were analyzed using the predictive tool and a score-based binning was performed to evaluate the cumulative antioxidant potential of the hydrolysate considering both predicted free radical scavenging (FRS) and metal chelating (CHE) peptides. As the hydrolysate is complex and peptides cannot be explicitly quantified, the raw peptide MS1 intensity was used as a proxy for peptide-level abundance as previously described (Bjørlie et al., 2024; Gregersen Echers et al., 2023; Jafarpour et al., 2020). Briefly, the sum of relative intensities for peptides within a scoring bin (based on AnOxPePred FRS and CHE scores) was computed relative to the sum of intensities for all peptides (following filtering of contaminants and false positive peptide IDs) to quantify the relative proportion of peptides with increasing confidence as antioxidants.

### 2.9. Statistical analysis

Analysis of variance (ANOVA) was conducted using SPSS Version 28.0.1.1 (Statistical Graphics Corp., Rockville, MD, USA), and means were compared pairwise using the Tukey test. The interfacial properties tests were performed twice independently, and the middle three cycles per amplitude were used for the rheological moduli. For analysis of specified protein families/groups, comparative analysis was performed as unpaired t-tests in GraphPad Prism (v.10.0.2, build 232). Comparisons of means were performed with using Welsh’s correction at 95% confidence intervals assuming Gaussian distributions and two-tailed p values.

## 3. Results and discussions

### 3.1. Crude protein content and degree of hydrolysis

According to the protein content results based on Dumas (Table S1), DC demonstrated higher crude protein (CP) content than DCH. This observation may be due to removal of some insoluble peptides during centrifugation. Moreover, both DC and DCH showed lower protein content than controls.

The degree of hydrolysis (DH) is essential in terms of techno-functional properties in protein hydrolysates. An optimized DH is essential for improving both solubility and functional properties without excessive peptide fragmentation, that could adversely influence techno-functionality. Using the conditions based on previous optimization work, (Badfar et al., 2026) hydrolysis of DC using 1% (w/w) Formea® Prime led to degree of hydrolysis (DH) of 10.98 % (Table S1), representing a moderate level of peptide bond cleavage. Hydrolysis for 2.75 h effectively disrupted the native DC protein structure, forming shorter peptides. Similarly, previous studies reported DH% close to 10% for trypsin-treated plant proteins such as pea and mulberry leaf protein, under comparable conditions (Shuai et al., 2022; Sun et al., 2021).

### 3.2. Interfacial properties

#### 3.2.1. Interfacial tension

Processing conditions significantly impact the surface activity and mechanical properties of the proteins at the interfacial layer. (Zhou et al., 2021). The amphiphilic nature of proteins enables them to adsorb at interfaces and form a viscoelastic network, aiding the interface to stabilize thermodynamically (Badfar et al., 2025). Interfacial tension (IFT) analysis was carried out on CGP and control samples using 0.1 % (w/w) protein concentration and monitored over 30 min (Fig. 1A). Initially, the instrument calibration was assessed by measuring interfacial tension behavior of water droplets in the MCT oil phase. The water droplets recorded a very constant response (≈26 mN/m), demonstrating the system’s calibration and the existence of no surface-active components. A similar interfacial tension value of water droplet in MCT oil was reported in a previous study (Badfar et al., 2026). Plant proteins can decrease interfacial tension and form viscoelastic interfaces, though they are not as soluble as animal protein sources, such as caseinate and albumins (Wojciechowski, 2022). Accordingly, Na-Cas, as a well-established animal protein source, could immediately be absorbed into the oil-water interface, rearrange conformation, and reduce IFT (12.75-11.79 mN/m). However, the plant protein samples (DC, DCH, PPI, and SPI) displayed slightly higher IFT values. Among them, the DC protein displayed a similar pattern of decreasing as Na-Cas and rapidly adsorbed at the oil interface, nevertheless with the higher values (15.52-12.48 mN/m). Similarly to an earlier study (Badfar et al., 2026), the DC hydrolysate (DCH) demonstrated a better ability to decrease IFT at equilibrium (≈13 mN/m) than the non-hydrolyzed DC (≈14 mN/m). This result revealed that the enzymatic hydrolysis facilitated increased IFT reduction by decreasing protein size to smaller peptide fragments and subjecting natively buried hydrophobic residues to the water-oil interface and potentially producing more amphiphilicity (Pérez-Gálvez et al., 2024). The PPI and SPI controls reached lower IFT values than the CGPs after 30 min. The IFT value for SPI was shown to be (≈12 mN/m) after 30 min, which is in line with what was reported in Wan et al. (2014). The result of PPIs IFT agrees with previous studies (Badfar et al., 2026). The effect on interfacial tension by proteins is highly related to not only the source of the proteins but also found to vary in proteins obtained from the same plant source using different processing techniques (Fig. 1A). These differences might be related to the amino acid composition of each sample, protein- (or peptide)-level conformation, and the rate and ability to undergo structural rearrangement at the water-oil interfaces, which can finally affect the surface activity of the proteins. To gain a more comprehensive understanding of the molecular behavior of different proteins at the interfaces, the dilatational rheological properties were studied.

**Figure 1.**
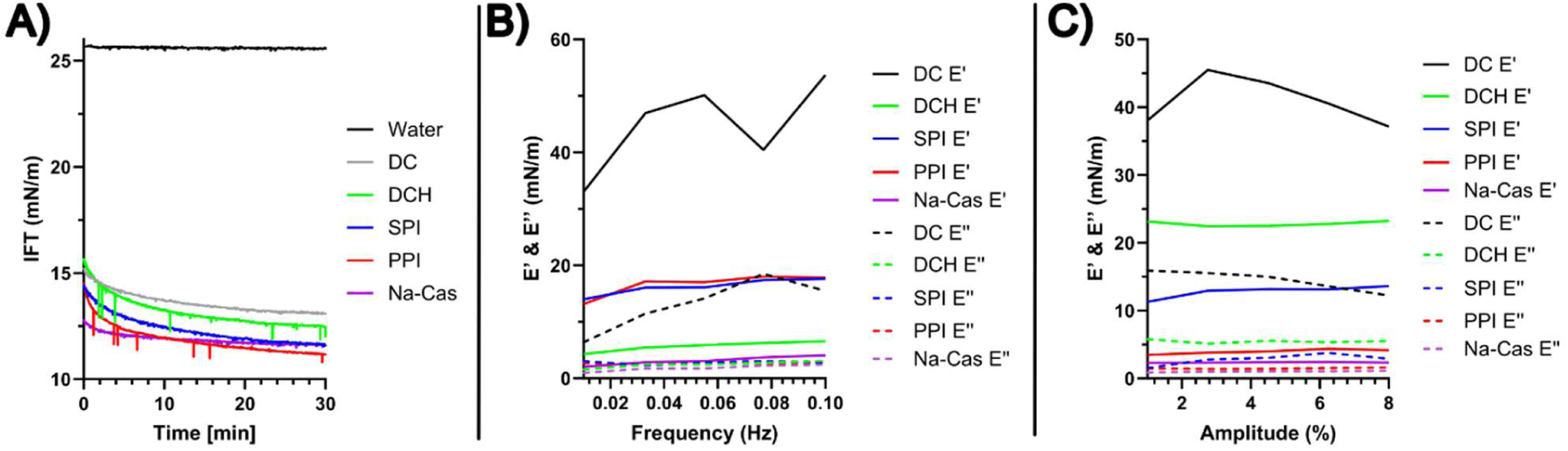
Interfacial tension reduction and dilatational rheology of selected clover grass protein protypes in comparison with a commercial pea protein isolate (PPI) soy protein isolate (SPI) and sodium caseinate (Na-Cas). A) Interfacial tension by pendant drop (water in MCT oil) using 0.1% (w/v) protein solutions over 30 minutes. B) Elastic (E’) and viscous (E’’) moduli from the frequency sweep test from 0.01 Hz to 0.1 Hz. C) Elastic (E’) and viscous (E’’) moduli from the amplitude sweep test from 1& to 8% amplitude.

#### 3.2.2. Dilatational rheology

A frequency sweep test was performed on CGPs and control samples within a range of 0.01-0.1 Hz at 4% amplitude (Fig. 1B). The results showed that E’ (the elastic modulus) was above E’’ (the loss modulus) in all samples. This relationship is characteristic of viscoelastic interfaces (Badfar et al., 2022). Except for the DC sample, all samples exhibited minimal dependency on the frequency range changes. At lower frequencies, increasing the frequency led to an increase in the E’ in the DC sample, indicating a more elastic response and resistance to deformation, as previously demonstrated (Badfar et al., 2026). In the frequency range of 0.06-0.08, the elastic modulus decreased, possibly due to the disruption of the interfacial structure and the inability of the DC proteins to reorganize rapidly with increasing frequency. However, with further frequency increase, the E’ increased again, suggesting the formation of a rigid and more elastic interface. This pattern reveals that the oil-water interface formed by the DC sample exhibited a complex and frequency-dependent viscoelastic response, likely due to the molecular interactions and physical properties of the DC protein at the interface. A higher elastic modulus has been suggested to indicate a higher rigidity of the interface (Tamm & Drusch, 2017). Among the plant proteins investigated, DC formed the most rigid and the least stretchable interface, followed by PPI, SPI, and DCH. Na-CAS displayed the lowest E’ and thus a higher stretchability than all plant samples, similarly to our previous findings (Badfar et al., 2026). These findings illustrate that scaling of the membrane-based biorefinery concept retains similar interfacial properties of the obtained CGPs.

Next, an amplitude sweep test was carried out on the CGPs and controls in the range of 1 to 8% deformation at a constant frequency (Fig 1C). The amplitude sweep test provides a valuable description of the dynamics of dispersions. The results reveal that applying the range of deformation displayed different patterns in each sample. For DC, increasing the amplitude from 1% to 3%, resulted in an increase in E’ (from 38.00 to 43.57 mN/m), indicating the formation of a more rigid or elastic interface (Yesiltas et al., 2023). This behavior suggests that DC protein at the interface shapes a structured network, facilitated by protein unfolding and structural rearrangement (Félix et al., 2019). In a previous study (Badfar et al., 2026), under the same condition and amplitude range, the DC produced in smaller scale showed a different response and by increasing amplitude from 1% to 3%, E’ deceased (from 15.40 to 15.10 mN/m). These comparisons show that DC in the current study formed much rigid and structured network than previous study, which may be attributed to differences in biomass composition, seasonal variation, or a direct result of process scaling.

At higher amplitude, the storage modulus started to diminish, and the interface weakened. This gradual reduction of E’ suggests that the protein network has been disrupted under higher amplitude. In contrast, DCH displayed a different pattern as both the storage modulus and the loss modulus in DCH were lower than in DC, indicating a less elastic interface, as previously demonstrated (Badfar et al., 2026). The constant storage modulus (E’) at lower amplitudes (1-3%) indicates the formation of a stable interfacial structure. When the amplitude increased, both the E’ and E’’ increased slightly, suggesting that the interface tends to deform with interfacial interactions gradually losing their stability under higher amplitudes. However, the interface remains viscoelastic since the storage modulus is higher than the loss modulus (Badfar et al., 2022). For SPI, E’ constantly increased from 1 % to 8 % amplitude compared to both CGPs. This initial increase was faster, indicating formation of a viscoelastic interfacial layer. At amplitudes higher than 3%, E’ increased slightly, suggesting a transition into a more stable interfacial layer, as the pattern showed that the interfacial behavior is less responsive but still adjusts and rearranges the molecules. The Na-Cas and PPI samples displayed an almost constant E’ and E’’, indicating a linear viscoelastic regime, and a stable interfacial layer (Yang et al., 2020). This observation reflects a stretchable but weaker interface structure (García-Moreno et al., 2021; Yesiltas et al., 2023). Since the two E’ and E’’ moduli have values close to each other, this suggests a transition point where further increases in the strain level can dominate the fluid or viscous dynamics. These observations suggest that protein composition and molecular interactions influence the dynamic behavior of each formed interfacial layer.

### 3.3. Physical stability

#### 3.3.1. ζ-potential, droplet size, and visual analysis

The ζ-potential of emulsions stabilized by CGPs and controls were determined to estimate electrostatic repulsions between fish oil droplets in the emulsions (Table 1). Measuring the electrostatic repulsion is meaningful as it can impact both the physical and oxidative stability of emulsions (Sha et al., 2021). The ζ-potential of the emulsions ranged from −49.87 mV to −30.25 mV on day 1 and −50.62 mV to −28.55 mV on day 8. The negative surface charge represents the availability of electron-rich molecules adjacent to the surface of emulsions (Zhang et al., 2021). CGPs displayed significantly higher absolute values of surface charge compared to the plant controls (PPI and SPI) and the animal control (Na-Cas) (p<0.05). A similar finding was found in a previous study, which showed that electrostatic repulsion between droplets in a rapeseed oil emulsion system, stabilized by different CGP prototypes, was significantly stronger than in PPI and Na-Cas emulsions (Badfar et al., 2026). The high negative potential for emulsions stabilized by CGPs suggest that they could discourage aggregation and maintain resistance due to heightened repulsion interaction between fish oil droplets. Moreover, it means that in the emulsion stabilized by CGPs, more ionic groups per unit surface area are presented than Na-Cas, SPI, and PPI. However, the stability of an emulsion is also a function of the emulsion’s particle size, which may result in a different distribution of hydrophobic and hydrophilic proteins and subsequently lead to different interfaces and different emulsion stabilities.

**Table 1.**
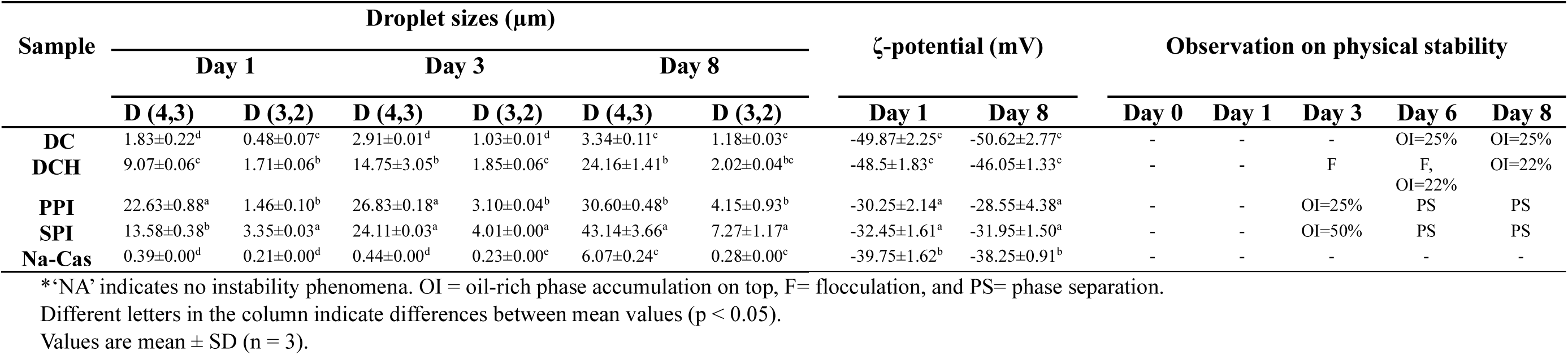
Values of Droplet sizes, ζ-potential and observation for physical stability of 5% fish oil emulsions produced using 0.4% (w/w, protein basis) clover grass (DC and DCH), pea isolate (PPI), soy isolate (SPI) and sodium caseinate (Na-Cas) protein solutions.

To investigate this further, the surface-weighted (D[3,2]) and volume-weighted (D[4,3]) mean diameter of droplets in the emulsions was analyzed over eight days of storage (Table 1). The type of emulsifier and homogenization can affect the size of the emulsion droplets (McClements et al., 2022). Both D[4,3] and D[3,2] of all emulsions increased from day 1 to 8 of storage. DC and Na-Cas achieved the lowest D[4,3] and D[3,2] among all samples, suggesting that DC and Na-Cas had faster diffusion to and adsorption at the droplet surface. However, the Na-Cas emulsion demonstrated a rapid increase in D[4,3] from day 3 to 8, but D[3,2] remained stable. Similar results have been reported previously for Na-Cas by (Badfar et al., 2025). Na-Cas, as an animal source of protein, is an adaptive protein lacking covalent sulfur bonds and can readily alter its conformation during the adsorption process and promote the formation of smaller droplets (Yesiltas et al., 2019).

Moreover, DCH formed smaller droplets (both D[4,3] and D[3,2]) compared to the plant controls (PPI and SPI). In a previous study, it was reported that at lower protein concentrations, RuBisCO-rich protein ingredients and soy protein are not able to form small droplets compared to whey protein, suggesting that some proteins may not adsorb at the oil droplet rapidly enough during the homogenization process (Tan et al., 2022). In previous work (Badfar et al., 2026), we found that DC produced at smaller scale could produce smaller droplets (D[4,3] and D[3,2]) compared to its hydrolysates, showing the opposite result from the current study. These discrepancies are likely due to the differences in biomass composition of CGPs in these studies and/or seasonal variation. Furthermore, the prototype in the current work comes from an upscaled version of the process. Process scaling may have an effect of properties of the product as well. Therefore, when the emulsifier does not sufficiently cover the oil droplets, coalescence occurs and facilitates the formation of larger oil droplets (Tan et al., 2022). Hence, achieving smaller droplets by DCH, PPI, and SPI may be possible by increasing the concentration of protein in the emulsion solutions.

Visual assessment of emulsion appearance was performed during the storage period (Table 1). One of the most common phenomena in emulsion instability is creaming, which can be seen by the migration of oil droplets into an upper phase separate from the remaining transparent water phase in the bottom. All emulsions were stable on day 0 and day 1. Subsequently, PPI and SPI emulsions displayed accumulation of an upper oil-rich phase (25 and 22%, respectively) on day 3. The oil-rich phase accumulation on top of an emulsion is similar to, but distinct from, creaming and is conventionally described as the oil index (OI). The phase at the bottom of oil-rich accumulation is not as transparent as in the phase at the bottom of creaming phenomenon (Yesiltas et al., 2021). The OI phenomenon may be due to the slow adsorption of protein molecules at the interface and subsequent oil droplet coalescence before protein has anchored at the oil droplet surface. This leads to larger oil droplets and the migration of the resulting droplets to the top of the emulsion (Yesiltas et al., 2021). On day 8, both emulsions stabilized with SPI and PPI displayed complete phase separation, which indicates a high degree of emulsion instability. This aligns with the results of ζ-potential and droplet sizes, which showed the lowest absolute value of repulsion between droplets and the larger droplet formation among all samples. For CGP emulsions, the DCH emulsion started to flocculate and accumulate an oil phase at the top of the emulsion from day 3. In contrast, the emulsion stabilized with intact and native CGP (DC) showed a stable system on day 3 without observed flocculation. Thus, the protein hydrolysate, DCH, may have demonstrated a lower physical stability than the intact protein, DC. This could be due to the existence of peptides with short chains and low molecular weight, prone to create an interfacial film with a lower viscoelasticity and cohesion (Hinnenkamp & Ismail, 2021). Although the interfacial measurement showed that DCH was able to reduce IFT more than DC after 30 min, the visual assessment exhibited no relationship between lower IFT value and physical stability. Similar observations have previously been reported in studies of the physical stability of emulsions (Berton-Carabin et al., 2018; Yesiltas et al., 2021).

### 3.4. Oxidative stability

#### 3.4.1. Peroxide value (PV)

Emulsions stabilized with CGPs (DC and DCH) exhibited the highest ability to protect the fish oil content from producing hydroperoxides during 9 days of storage (Fig 2). The initial PV of emulsions stabilized with PPI, SPI, and Na-Cas ranged from 4.31-7.51 meq O_2_/kg oil, increasing from day 0 to day 9. The high PV at day 0 could indicate oxidation during the emulsion production process, as the high shear stress and incorporation of oxygen may happen (García-Moreno et al., 2016; Horn et al., 2012). Based on statistical analysis (Table S2), the PV value did not increase significantly for any emulsion within the initial phase of the storage, indicating a lag phase in the oxidation processes. However, the length of the lag phase varied between emulsions, where the Na-Cas emulsions showed the longest lag phase (day 0 to day 5) and SPI showed the shortest lag phase (day 0 to day 1). The remaining emulsions (PPI, DC, and DCH) all showed an intermediate lag phase (day 0 to day 2). The high PV value observed for the Na-Cas emulsion on day 9 is higher than values reported in previous studies. Ballon et al., (2025) reported a peroxide value (PV) of approximately 27 meq O₂/kg oil after 9 days of storage for a 5% fish oil-in-water emulsion stabilized with 0.2% sodium caseinate. Similarly, Queiroz et al., (2021) observed a comparable PV (27.33 meq O₂/kg oil) after 10 days of storage in a 5% fish oil-in-water emulsion stabilized with the same sodium caseinate concentration. The discrepancy could be attributed to different parameters, such as storage conditions, temperature, oxygen exposure, light, droplet size distribution, or an increased Na-Cas concentration. The latter could indicate a potential prooxidant effect of Na-Cas at higher concentrations affecting hydrogen peroxide formation and decomposition, but this warrants further investigation. The DC emulsion demonstrated the lowest formation of hydroperoxides until the last day of storage, followed by DCH. Moreover, CGP emulsions displayed a lag phase prior to hydroperoxide formation. The lower oxidation rate of emulsions stabilized with CGPs could potentially be ascribed to their ability to chelate transition metal ions at the interface and in the water phase or to an inherent scavenging of radicals.

**Figure 2.**
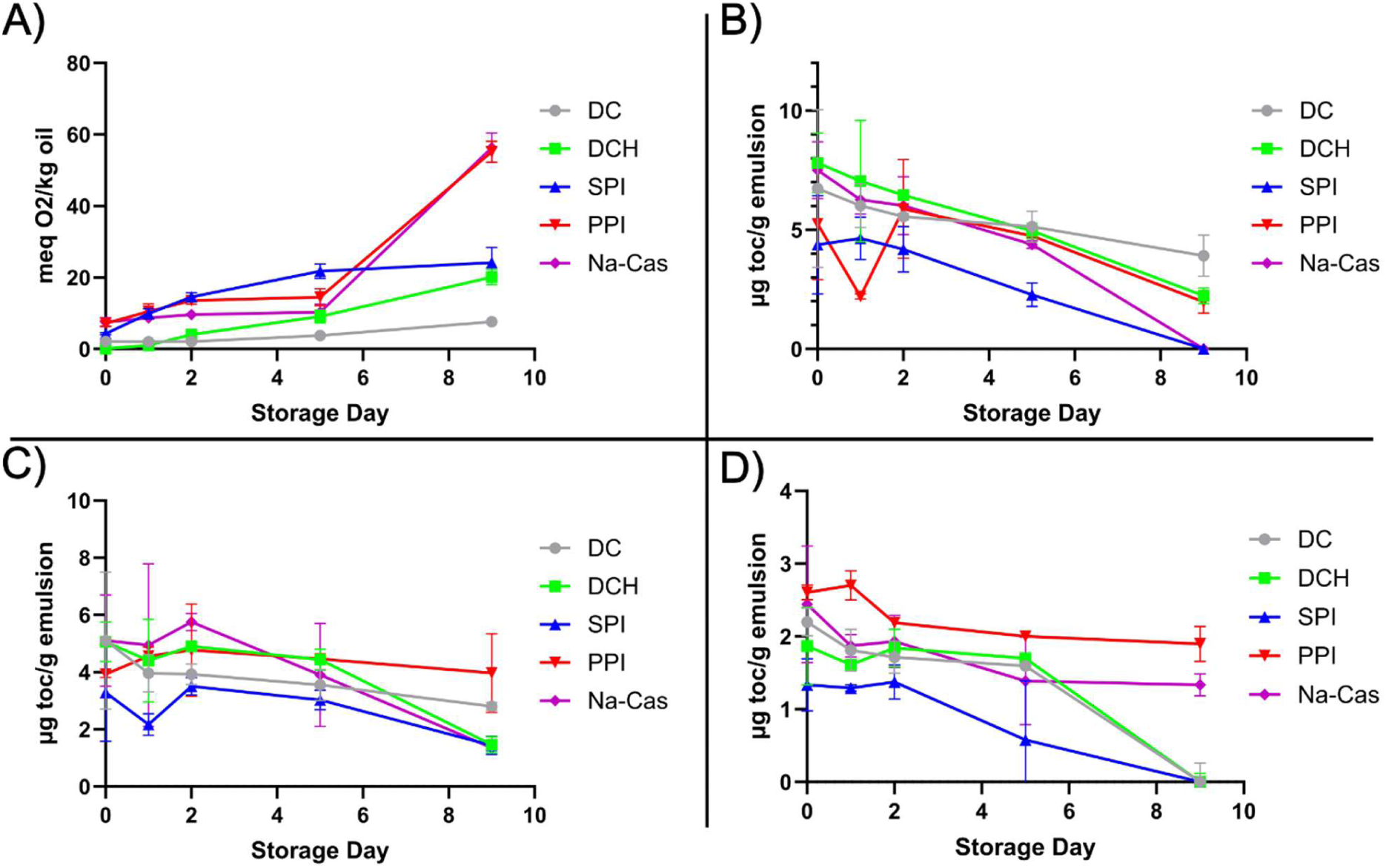
Peroxide value (PV) and tocopherol consumption in 5 % fish oil-in-water emulsions stabilized with Na-Cas, PPI, SPI, DC, and DCH (0.4 % w/w, protein basis) during 9 days of storage. A) Development of PV (meq O_2_/kg oil) during storage. Development of α- (B), γ- (C), and δ- (D) tocopherol content (µg toc/g emulsion) during storage.

#### 3.4.2. Tocopherols

Tocopherols are renowned natural antioxidants, available in their α-, β-, γ-, and δ–forms. They are lipid-soluble and naturally found in fish oil, with the exception of β-tocopherol, which is usually not present (Hu & Jacobsen, 2016). The mechanism of antioxidant activity for tocopherols is principally donating hydrogens to free radical lipids, thereby quenching oxidation propagation. The tocopherol content of the fish oil used for emulsions was determined to be 470.8, 345.5, and 211.0 (µg toc/g oil) for α, γ-, and δ—tocopherol, respectively. Tocopherol consumption in emulsions was monitored for 9 days of storage (Fig. 2B-D). β-tocopherol content is typically not included as the level in fish oil-in-water emulsions is too low (Irankunda et al., 2024). Results indicated that not only was the tocopherol content in the emulsions prepared with 5% fish oil significantly lower than expected for a 5 % oil mixture, but also their content decreased significantly during the nine storage days. Furthermore, the reduction in the tocopherol level at day 0 compared to that of fresh oil is evident of oxidation during the emulsification process. Significant differences between the emulsions observed already on day 0, indicate that not only were oxidative processes initiated during emulsion preparation but that the different protein emulsifiers displayed different levels of oxidative protection during production. The α-tocopherol content gradually decreased from day 0 to day 9 in all emulsions (Fig. 2B), which indicates that the tocopherol is consumed when preventing formation of primary and secondary oxidation products (Irankunda et al., 2024). However, this cannot be concluded based on tocopherol levels only. The reduction of α-tocopherol in emulsions stabilized with PPI and CGPs (DC and DCH) was slower. On day 9, α-tocopherol in Na-Cas and SPI emulsions was fully depleted and was significantly lower (p < 0.05) than the rest of the emulsions. The significant reduction in the level of α-tocopherol during the storage period can be attributed to the lack of a lag phase in peroxide formation (Bjørlie et al., 2023). Generally, tocopherol levels reduced during storage in DC and DCH. When considering the absolute reductions, α-tocopherol indicated the largest decrease, consistent with it being the predominant tocopherol initially present. Shifts in γ- and δ-tocopherol were smaller in absolute terms. Overall, PPI and CGPs were the most efficient in terms of preventing emulsion oxidation.

#### 3.4.3. Development of secondary oxidation products

Volatile secondary oxidation compounds, such as ketones, aldehydes, and alcohols, are formed during decomposition of primary oxidation products (hydroperoxides). Due to the high number of double bonds in long chain ω-3 PUFAs in fish oil, many different volatile compounds can be generated, leading to lower quality and off-flavor. Among the identified volatile compounds, (t,t)-2,4-heptadienal, and 1-penten-3-ol (Fig. 3) are typically the most abundant and odor active volatile compounds in oxidative deterioration of emulsions, and hence broadly applied to estimate the oxidative stability (Irankunda et al., 2024; Yesiltas et al., 2019, 2022, 2023). (t,t)-2,4-heptadienal and 1-penten-3-ol are derived from the oxidation of ω-3 fatty acids(García-Moreno et al., 2021). The rest of the detected compounds, including c-4-heptenal, octanal, t-2-hexenal, t-2-pentenal, 1-penten-3-one, pyridine, t-2-butenal, butanal, pentanal, 1-pentanol, and hexanal are available in the supplementary file (Fig. S1).

**Figure 3.**
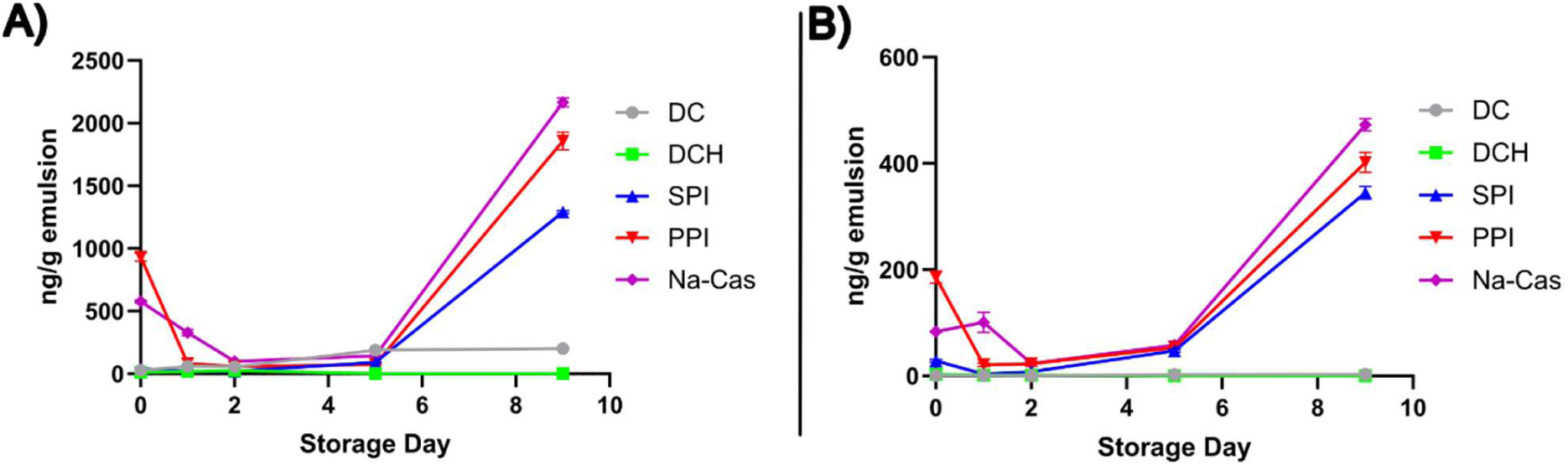
Formation of volatile secondary oxidation products. Quantification 1-penten-3-ol (A), and t,t-2,4-heptadienal (B) in 5 % fish oil-in-water emulsions stabilized with 0.4% (w/w, protein basis) Na-Cas, PPI, SPI, DC, and DCH over 9 days of storage.

Plant protein (SPI and PPI) and animal protein (Na-Cas) controls were found to not prevent the fish oil from further oxidation (Fig. 3) in agreement with the peroxide value (PV) and tocopherol consumption results (Fig. 2). Levels for 1-penten-3-ol and 2,4-heptadienal (as well as butanal, octanal, and pentanal (Fig. S1)) until day 5 were in line studies by Queiroz et al., (2021) and Ballon et al., (2025), although these studies found lower concentrations on day 9 and 10. Differences in volatile compound levels on day 9 in comparison with previous studies may partly be attributed to variations in sample handling prior to storage. In particular, insufficient nitrogen flushing of sample bottles on day 9 could have resulted in higher oxygen availability in the headspace, thereby promoting lipid oxidation even during frozen storage. Nevertheless, all samples on day 9 were treated identically, and CGP emulsions did not show any indications of volatile product formation.

The lower concentration of volatile secondary oxidation compounds in CGP emulsions is attributed to inhibition of lipid oxidation. In all analyzed volatile compounds, DC and DCH have very low or no formation during emulsion storage, indicating higher oxidative stability compared to PPI, SPI, and Na-Cas. It can be suggested that CGPs may function as strong metal chelators, thereby preventing the formation of volatile compounds through cation-induced oxidation. The presence of (*t,t*)-2,4-heptadienal has been associated with nasty, fishy, and rancid odors in emulsions, while 1-penten-3-ol causes a sweet odor (Hartvigsen et al., 2000). DC and DCH emulsions did not exhibit these products during 9 days of storage based on GC-MS analysis. Overall, the results suggest that, compared to SPI, PPI, and Na-Cas, emulsions with CGPs exhibit a greater inherent oxidative stability as CGPs effectively inhibit the formation of primary and secondary oxidation products in the oil-in-water emulsion system, regardless of whether enzymatic hydrolysis is applied or not. Although GC–MS analysis provides valuable information on the formation of volatile oxidation products, the final odor perception of the samples cannot be concluded from instrumental data alone. Odor perception is influenced by the odor thresholds of individual compounds, which are matrix dependent. In addition, interactions among multiple volatile compounds may result in synergistic or masking effects. Therefore, sensory evaluation would be required to directly assess the perceived odor of the samples.

### 3.5. Protein- and peptide-level characterization of CGPs

To investigate why GPCs were able to reduce lipid oxidation in the emulsions, a proteomic analysis was performed. The analysis included the two GPCs as well as the initial, unfiltered green juice (Feed) used to produce DC via the two-stage membrane filtration approach (Mattsson et al., 2025). Across all samples, a total of 1921 proteins were identified following filtering of false positives and contaminant proteins (Table S3). In the initial feed solution, a total of 1548 proteins were reproducibly quantified (i.e. intensity values were recorded in at least two of three replicates). In the concentrate (DC), 963 proteins were reproducibly identified while in DCH, peptides from merely 69 proteins were reproducibly quantified. Qualitatively, the overlap of reproducibly observed proteins shows that among the 1786 reproducibly identified proteins in any sample, only 63 (3.5 %) were reproducibly observed in all three streams (Fig. 4A). This can be ascribed to the low number of IDs in DCH, as 662 (37 %) proteins were reproducibly identified in both Feed and DC. A substantial proportion (821 proteins, 46 %) were only observed in the initial feed, while a smaller proportion (238 proteins, 13 %) were only observed in one or both downstream samples (DC and DCH). That some proteins were only identified in downstream samples, particularly DC, can be ascribed to a reduction in sample complexity going from the initial feed to the final concentrate over the two-stage filtration process and subsequent diafiltration. Reducing sample complexity improves detectability of low abundant proteins, in line with previous studies on small scale production of DC (Badfar et al., 2026; Gregersen Echers et al., 2026). Among the feed-exclusive proteins, a range of chlorophyll-binding proteins (CBPs) and other pigment-binding proteins from photosystems I and II were observed. These observations agree with previous studies on a similar two-stage membrane filtration process at smaller scale, where the first-stage filtration selectively retained these proteins, thereby eliminating the vast majority of green pigments from the first stage permeate and subsequent streams (Badfar et al., 2026; Gregersen Echers et al., 2026).

Comparing Feed and DC using MaxLFQ data, clear differences in protein composition were observed (Fig. 5B). In the quantitative comparison using MaxLFQ data, DCH was excluded based on the low number of protein IDs (Fig. 5A), which may skew the interpretability from a comparative analysis.

**Figure 5:**
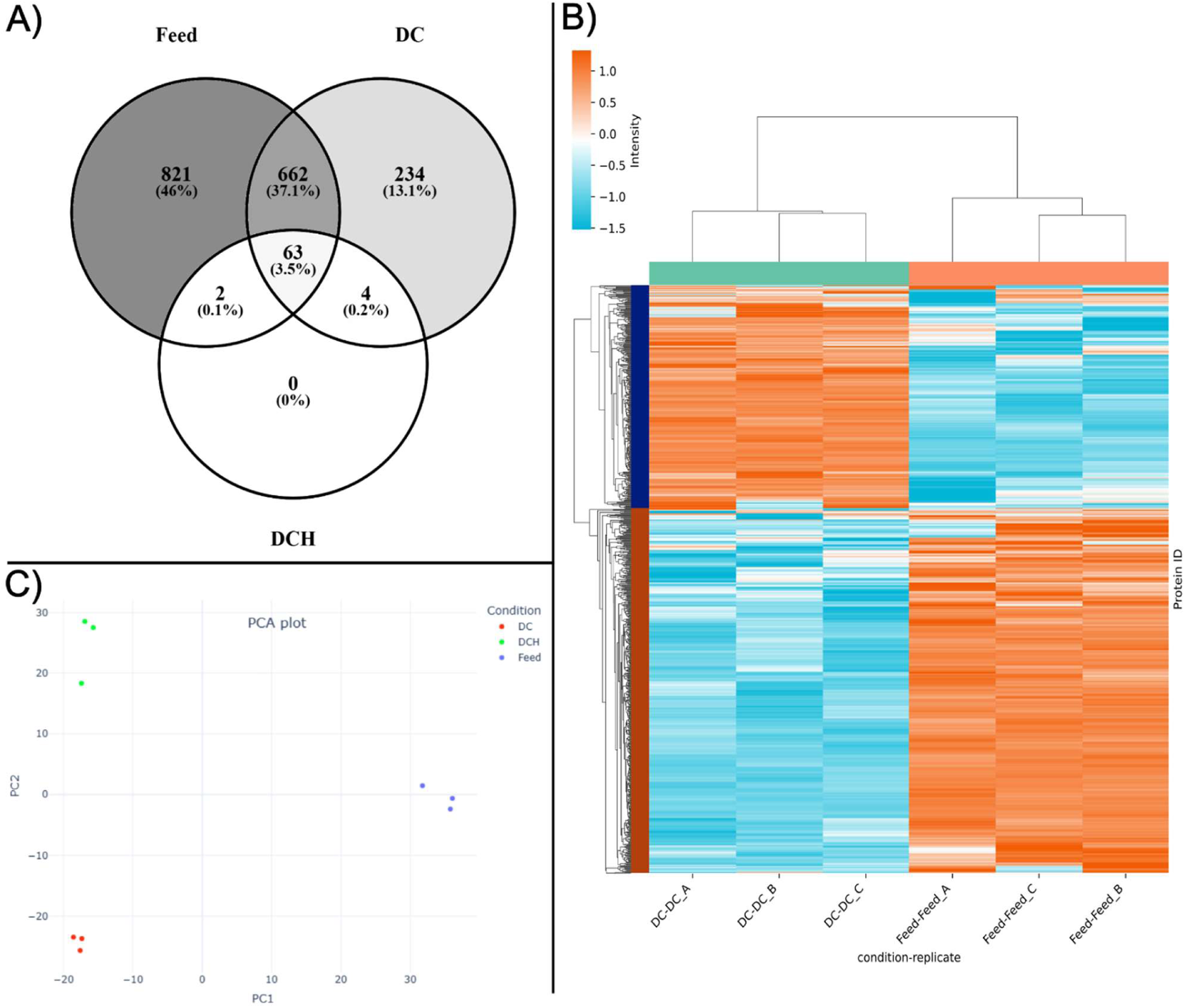
Comparative analysis of reproducibly quantified proteins and MaxLFQ data. A) Venn diagram of reproducibly quantified (i.e. quantified in at least two of three replicates) proteins across Feed (green juice), DC, and DCH showing the number and proportion (%) of protein IDs within the different samples and their overlap. B) Heatmap representation of differentially abundant proteins for the initial Feed and DC by ANOVA analysis in MassDynamics. Data is depicted as z-score standardized MaxLFQ intensities by row (protein group) and clustered using a Euclidian distance of 3. C) Two-dimensional representation of data variability by PCA analysis using MaxLFQ-based quantification for Feed, DC, and DCH.

The pair-wise analysis revealed that 435 proteins were enriched in Feed while 248 proteins were enriched in DC. Among the most enriched proteins in Feed, ATP synthase subunit α (ATPs-α) (A0A2Z6M310, log2FC = 9.1), ATP synthase subunit β (ATPs-β) (A0A023HPU4, log2FC = 9.0), Photosystem II D2 protein (PII-D2) (A0A023HQ62, log2FC = 7.9), and Chlorophyll a-b binding protein (CBP) (A0A2K3LTY7, log2FC = 7.6) were found. These observations both corroborate qualitative analysis and agree with previous studies also finding CBP, PII proteins, and ATPs depleted after a similar membrane filtration performed at smaller scale on different green biomasses (Badfar et al., 2026; Gregersen Echers et al., 2026) In DC, a different subset of clover grass proteins was found to be enriched. This subset included e.g. superoxide dismutase (SOD) (I1I9J4, log2FC = 6.2) and peroxidase (A0A2Z6P5L2, log2FC = 4.9) but also several endogenous proteases such as thiol protease aleurain-like protein (A0A2K3N629, log2FC = 5.7), putative serine protease EDA2-like protein (A0A2K3L3V8, log2FC = 5.7), and subtilisin-like protease-like protein (A0A2K3KYU8, log2FC = 5.6) among the most enriched proteins. While proteins with known antioxidant properties have also previously been found to be enriched following two-stage membrane filtration at smaller scale (Gregersen Echers et al., 2026), and may explain the improved oxidative stability of DC-based emulsions, the enrichment of endogenous proteases in the product stream (DC) also substantiates the need for process control to minimize uncontrolled hydrolysis during green juice processing into a protein ingredient (Badfar et al., 2026). The difference in both qualitative and quantitative composition between not only Feed and DC but also DCH is also reflected when visualizing inter-sample variability using PCA (Fig. 5C). Feed differentiates from DC and DCH primarily on the first PC, representing 31% of the total variability, while DC and DCH differentiates with the second PC, accounting for 21% of the variability within the dataset. While clear quantitative differences were found by MaxLFQ analysis, a complementary quantification using relative intensity-based absolute quantification (riBAQ) was performed to identify the most abundant proteins within the different samples.

Across the Feed and DC replicates, 48 different proteins were found to each account for at least 0.5 % of the protein by mean riBAQ) in either of the samples (Fig. 6A). Together, these 48 proteins represent 56% of the protein in Feed and 69% of the protein DC, but also 65% of the (inferred) protein in DCH (Table S4). Similarly to earlier studies (Badfar et al., 2026; Danner Aakjaer Pedersen et al., 2025; Gregersen Echers et al., 2026) the most abundant proteins were the large and small subunits of RuBisCO (rbcL and rbcS, respectively) across all samples, but also CBP and different photosystem proteins in the initial green juice Feed which, in turn, were depleted in both DC and DCH. In contrast, proteins such as chitinase, malate dehydrogenase, ferredoxin-NADP (+) reductase were found in substantially higher mean abundances in the downstream DC and DCH samples. As several of these have previously been associated with *in vitro* endogenous antioxidant activity in green juice (Danner Aakjaer Pedersen et al., 2025), we determined the cumulative abundance of ten previously reported antioxidant proteins in a family-wise manner across the samples, in addition to the cumulative abundance of RuBisCO, CBPs, and photosystem I and II (PI/II) proteins (Fig. 6B). From the family-wise analysis, a significant enrichment of rbc in DC was found, while the cumulative abundance of both CBPs and PI/II proteins was significantly depleted from a substantial abundance in Feed (ΣriBAQ =16%) to a principally non-present level in DC (ΣriBAQ < 0.03%), representing a 99,8% removal. In contrast, known antioxidant proteins such as ferredoxin-NADP reductase (FNR), lactoylglutathione lyase (Glyoxalase I; Glo1), glutathione S-transferase (GST), peroxidase (Px), superoxide dismutase (SOD), and thioredoxin-dependent peroxiredoxin (TPx) were significantly enriched in DC compared to Feed. Taken together, the abundance of the ten protein families with previously reported antioxidant activity *in vitro*, increased from 2.3% in Feed to 9.8% in DC, representing a 4.3-fold enrichment following the membrane-based biorefinery process (Fig. 6B). That antioxidant proteins constitute 10% of the total protein in the native CGP concentrate can likely explain the improved oxidative stability of the DC-based emulsion compared to the PPI, SPI, and Na-Cas controls.

**Figure 6:**
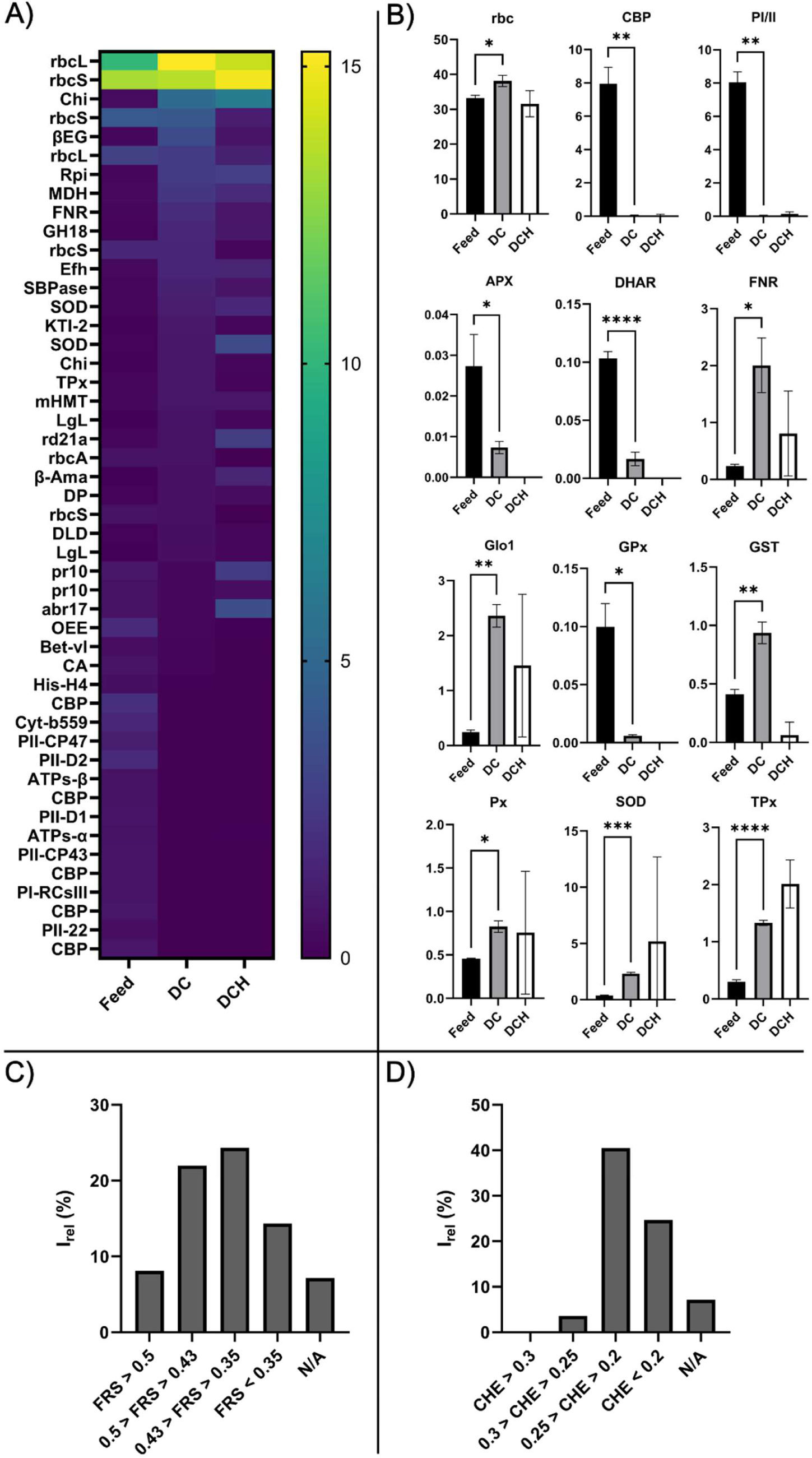
Relative protein and protein family distribution and predicted peptide-level antioxidant properties. A) Heatmap representation of relative molar abundance (mean riBAQ, %) distribution across Feed, DC, and DCH showing proteins of high abundance (riBAQ > 0.5%) in any sample. B) Cumulative abundance (sum of isoform riBAQ, %) within selected protein families/groups for Feed, DC, and DCH. Cumulative abundance is shown for RuBisCO (rbc), chlorophyll a-b-binding proteins (CBP), and photosystem I & II (PI/II) proteins as well as ten proteins families, previously reported to display potential endogenous antioxidant (AnOx) activity in their native forms (Danner Aakjaer Pedersen et al., 2025). AnOx proteins include: L-ascorbate peroxidase (APX), dehydroascorbate reductase (DHAR), ferredoxin-NADP reductase (FNR), lactoylglutathione lyase (Glyoxalase I; Glo1), Glutaredoxin-dependent peroxiredoxin (GPx), glutathione S-transferase (GST), Peroxidase (Px), Superoxide dismutase (SOD), and Thioredoxin-dependent peroxiredoxin (TPx). Comparison of means was performed using a double-tailed Welsh t-test with a confidence interval of 95%. Significant differences between means (from adjusted p-values) are shown as “ns” (p > 0.05), “*” (p ≤ 0.05), “**” (p ≤ 0.01), “***” (p ≤ 0.001), and “****” (p ≤ 0.0001). Due to large standard deviations in DCH due to the low number of peptide IDs, comparison of means was only performed between Feed and DC. Peroxiredoxin Q-like (ycf33) was not identified in neither DC nor DCH and hence not included in the plot. C) Distribution of relative peptide-level MS1 intensity for reproducibly quantified peptides in DCH based on predicted free radical scavenging (FRS) activity. D) Distribution of relative peptide-level MS1 intensity for reproducibly quantified peptides in DCH based on predicted metal chelating (CHE) activity. For FRS and CHE, mean relative peptide MS1 intensities (I_rel_ (%)) are binned according to prediction scores using AnOxPePred (Olsen et al., 2020).

On the peptide-level, DCH was found to be of low complexity. The number of total and reproducible peptide IDs in Feed was 4819 and 4050 while in DC, a total of 3092 peptides and 2626 reproducible peptides were identified (Table S5). This further substantiates the selective protein-level retention in the first stage filtration going from initial Feed to downstream samples (e.g. DC), as previously demonstrated (Badfar et al., 2026; Gregersen Echers et al., 2026). In contrast, across the triplicate analysis of DCH, merely 496 peptides were identified and only 89 of these were reproducible. The low number of peptide IDs in DCH indicates selectivity during hydrolysis as well as loss of protein. While the content of peptides originating from the main protein constituent (RuBisCO) in DCH reflects a similar level as in Feed and DC (Fig. 6A-B), many other abundant proteins (i.e. their peptides after tryptic digestion) were not identified in DCH. This agrees with Linderstrøm-Lang theory, stating that preferential hydrolysis will occur and that during *in vitro* hydrolysis, a protease is more likely to “finish” hydrolysis of one individual protein before moving to the next (Christensen et al., 2025). As such, complete hydrolysis, as obtained under highly optimized conditions as those used under proteomics sample preparation (DC), is not comparable to those observed during partial *in vitro* hydrolysis with subsequent heat inactivation (DCH).

Nevertheless, the identified peptides in DCH were analyzed *in silico* using AnOxPePred (Olsen et al., 2020) to investigate if the improved oxidative stability seen in emulsions could potentially be attributed to peptides with antioxidant potential (Table S6). The 89 reproducible peptides represent 76 % of the average, relative peptide intensity across the triplicates, thereby explaining the vast majority of the measured signals. Based on previous studies using AnOxPePred, a score > 0.43 for free radical scavengers (FRS) represents a peptide with higher probability of displaying *in vitro* FRS activity than not (Bjørlie et al., 2024; Jafarpour et al., 2020). Similarly, a cutoff score of 0.30 applies for metal chelating peptides (CHE). For FRS scores, 30 % of the total signal originates from peptides with FRS scores > 0.43 and 8.1 % from peptides with FRS scores > 0.5 (Fig. 6C). Among the highest scoring and most abundant (by relative MS1 intensity) peptides, EHNNSPGYYDGR (FRS = 0.54) from *Trifolium pratense* rbcS (A0A2K3P1N8) constitute 3.9% of the average peptide MS1 signal while LSGGDHIHAGTVVGK (FRS = 0.48) from *Trifolium repens* rbcL (U6BRQ4) constitutes 14.1%. These observations underline that while RuBisCO itself has only to a limited extent been linked with endogenous antioxidant activity by Mg^+^ chelation (Bathellier et al., 2020; Danner Aakjaer Pedersen et al., 2025), embedded peptides in both subunits are predicted to be potential radical scavengers *in vitro*. In contrast, no peptides were found with CHE scores above the 0.30 threshold among the 89 reproducibly identified peptides (Fig. 6D). As such, the observed retardation of lipid oxidation observed in DCH emulsions is likely attributed to radical scavenging activity.

## 4 Conclusion

In this study, two clover grass protein (CGP) samples (a filtration concentrate (DC) and its tryptic hydrolysate (DCH)) were investigated as protein-based emulsifiers. The dynamic behavior CGPs at the water-oil interface was analyzed using interfacial tension (IFT) and dilatational rheology using MCT oil in a comparative evaluation with commercial plant-(pea protein isolate (PPI), soy protein isolate (SPI)) and animal-based (sodium caseinate (Na-Cas)) protein ingredients. All plant proteins (GCPs, PPI, and SPI) exhibited higher interfacial tension values (15.52-12.48 mN/m) compared to Na-Cas (11.79 mN/m). Differences in rheological behavior between the sample suggested that protein composition and molecular interactions influence the dynamics of the formed interfaces. In 5% fish oil-in-water emulsions (0.2% protein (w/w)), both DC and DCH produced smaller droplet sizes and larger absolute electrostatic repulsion between droplets than the commercial plant protein controls (PPI and SPI) during the 8-day storage period (p < 0.05), indicating that CGPs provided better physical stability. These results were confirmed with visual analysis, as PPI and SPI underwent complete phase separation by day 6. CGP emulsions demonstrated superior resistance to oxidation by producing lower levels of both hydroperoxides as the primary oxidation product and reduced formation of volatile compounds as secondary oxidation products compared to SPI, PPI, and Na-Cas. Moreover, PPI and CGPs showed lower consumption of α-tocopherols, during storage, indicating higher oxidative stability of the CGP emulsions.

Based on quantitative proteomics analysis, the most abundant protein family in GPCs was not surprisingly RuBisCO, constituting almost 40% of the total protein in DC. Moreover, proteins with previously reported *in vitro* antioxidant activity constituted 10% of the total protein in DC, which explains the improved oxidative stability of DC-based emulsions. Based on bioinformatic analysis of reproducible identified peptides in DCH, 30% of the hydrolysate (based on relative peptide MS1 intensity) was constituted by peptides with probable free radical scavenging activity, while none of the peptides obtained a score above the threshold for probable metal chelating activity. As such, the ability of DCH to improve the oxidative stability of emulsions is likely linked to the high abundance of peptides able to scavenge induced radicals, thereby decreasing the rate of peroxide formation and lipid retardation. Overall, this study provides valuable insights into the interfacial and emulsifying properties of protein ingredients derived clover grasses. Moreover, this study improves our understanding of the molecular composition of such ingredients and link it to beneficial properties such as improvement of oxidative stability of emulsions. Although clover grass proteins are promising as a sustainable protein source for food technology, further studies are needed to improve understanding of molecular characteristics and suitability as a food protein ingredient for human consumption.

## 6. Acknowledgements

The authors of this paper would like to express their special thanks to lab technician Inge Holmberg from the Research Group for Bioactives-Analysis and Application, DTU, for her skillful assistance. Also, a special thanks to BiomassProtein A/S, and particularly site Manager Morten Olsen, for assistance with biomass harvesting, pressing, and filtration).

## 8. Author Contribution

NB: Methodology, Validation, Formal analysis, Investigation, Writing – original draft preparation, Writing – review and editing, Visualization. CJ: Resources, Writing – review and editing, Supervision. ML: Conceptualization, Methodology, Funding acquisition, Writing – Review & Editing, Supervision. SGE: Conceptualization, Methodology, Validation, Formal analysis, Investigation, Writing – original draft preparation, Writing – review and editing, Visualization, Supervision.

