## Supplementary information for "Protein concentrates from pilot-scale biorefining of clover grass: Interfacial properties and their effects on the physical and oxidative stability of fish oil-in-water emulsion"

**For**

### Content of supplementary material

#### Supplementary tables

**Table S1**. Protein content of clover grass proteins (CGPs) and controls and degree of hydrolysis for the DC hydrolysate.

**Table S2:** Statistical analysis of peroxide values (PV) development for emulsions produced by CGPs and controls.

**Table S3:** MaxQuant output data and additional downstream processing hereof.

**Table S4:** Overview of the most abundant (riBAQ > 0.5%) proteins in any of the analyzed samples.

**Table S5:** MaxQuant peptide-level output data and additional downstream processing hereof.

**Table S6:** Overview of reproducibly quantified peptides in DCH along with the predicted antioxidant potential.

#### Supplementary figures

**Figure S1.** Formation of volatile secondary oxidation products in %5 fish oil-in-water emulsions

**Table S1**. Protein content of clover grass proteins (CGPs) and controls and degree of hydrolysis for the DC hydrolysate. Values are indicated as mean ± SD (n = 3).

| Sample | Sample type | Crude Protein (%) | DH (%) |
|---|---|---|---|
| DC | Dia concentrate | 49.78±4.46 | N/A |
| DCH | Dia concentrate hydrolysate | 41.99±1.42 | 10.78±1.13 |
| PPI | Pea protein isolate | 87.00±0.00 | N/A |
| SPI | Soy protein isolate | 85.00±0.00 | N/A |
| Na-Cas | Sodium caseinate | 93.00±0.00 | N/A |

**Table S2:** Statistical analysis of peroxide values (PV) development over time for emulsions produced by CGPs and controls. Values are indicated as mean ± SD (n = 3). Different letters in the column indicate differences between mean values ($p < 0.05$).

| Day of Storage | DC | DCH | SPI | PPI | Na-Cas |
|---|---|---|---|---|---|
| 0 | 2,1 ± 0,42 $^{a}$ | 0,08 ± 0,02 $^{a}$ | 4,32 ± 0,29 $^{a}$ | 7,15 ± 0,74 $^{a}$ | 7,51 ± 1,3 $^{a}$ |
| 1 | 2,06 ± 0,54 $^{a}$ | 1,01 ± 0,08 $^{a}$ | 9,88 ± 1,39 $^{ab}$ | 10,5 ± 2,16 $^{a}$ | 8,73 ± 0,19 $^{a}$ |
| 2 | 2,11 ± 0,14 $^{a}$ | 4,02 ± 1,2 $^{ab}$ | 14,56 ± 1,21 $^{bc}$ | 13,55 ± 0,99 $^{a}$ | 9,61 ± 0,22 $^{a}$ |
| 5 | 3,78 ± 0,23 $^{b}$ | 9,07 ± 1,86 $^{b}$ | 21,76 ± 2,02 $^{cd}$ | 14,48 ± 2,34 $^{a}$ | 10,29 ± 2,25 $^{a}$ |
| 9 | 7,64 ± 0,1 $^{c}$ | 20,07 ± 2,13 $^{c}$ | 24,19 ± 4,3 $^{d}$ | 55,15 ± 2,93 $^{b}$ | 56,28 ± 4,14 $^{b}$ |

**Table S3:** MaxQuant output data and additional downstream processing hereof. The "proteinGroups" file has been modified to include a range of additional data used in the manuscript covering calculation of relative iBAQ (riBAQ), replicate means, abundance and reproducibility filters. For a full data file with no modification, see the referenced project in the PRIDE data repository. Table can be found in appended .xlsx file (Supplementary Table) as "Table S3".

**Table S4:** Overview of the most abundant (riBAQ > 0.5%) proteins in any of the analyzed samples. The table includes the Uniprot AC#, protein name, a short name (as used in Figure 4), various MaxQuant Data (number of proteins in group, peptide IDs, unique peptides, sequence coverage, molecular weight and Andromeda score), mean riBAQ abundance across all sample types. The table contains both individual proteins (MaxQuant proteinGroups) and the defined "families/types/groups" from this work. Table can be found in appended .xlsx file (Supplementary Table) as "Table S4".

**Table S5:** MaxQuant peptide-level output data and additional downstream processing hereof. The "peptides" " file has been modified to include a range of additional data used in the manuscript covering calculation of relative MS1 intensity ($I_{rel}$), replicate means, and reproducibility filters. For a full data file with no modification, see the referenced project in the PRIDE data repository. Table can be found in appended .xlsx file (Supplementary Table) as "Table S5".

**Table S6:** Overview of reproducibly quantified peptides in DCH along with the predicted antioxidant potential as free radical scavengers (FRS) and metal chelators (CHE) using AnOxPePred. Table can be found in appended .xlsx file (Supplementary Table) as "Table S6".

**Figure S1.** Formation of volatile secondary oxidation products in %5 fish oil-in-water emulsions stabilized with 0.4% (w/w, protein basis) Na-Cas, PPI, SPI, DC, and DCH over 9 days of storage.. Quantification (ng/g emulsion) of c-4-heptenal (A), octanal (B), t-2-hexenal (C), t-2-pentenal (D), 1-penten-3-on (E), pyridin (F), t-2-butenal (G), butanal (H), pentanal (I), 1-pentanol (J), and hexanal (K).

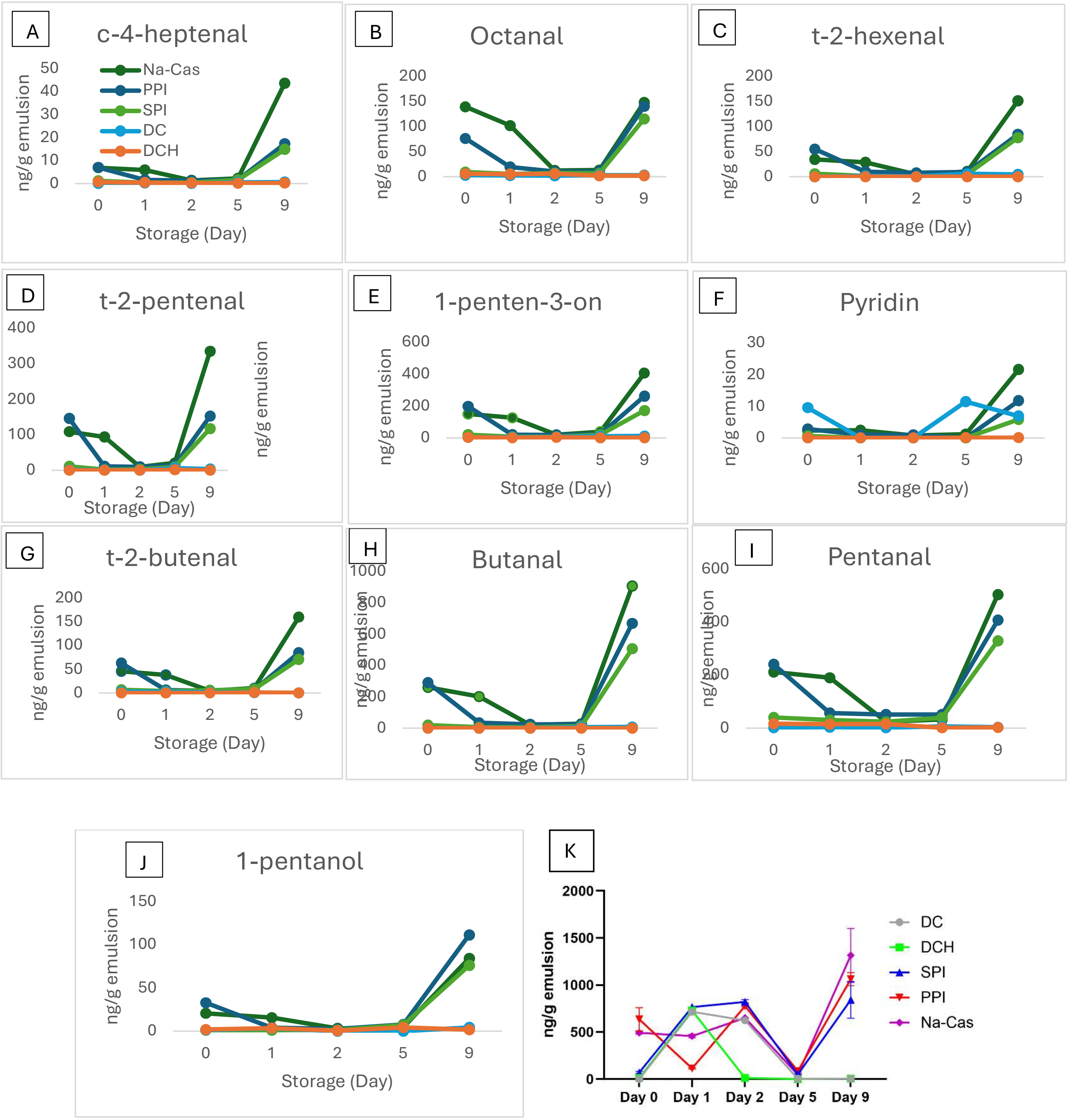
